## supplemental text for "Molecular tuning of sea anemone stinging" (from another version of this preprint: version 3, 2023-09-05)

### Supplemental Information

In the main text, we have introduced Markov Decision Processes that model the problems of defensive *vs* predatory stinging. In the following, we will show an approximate analytical solution to Bellman's optimal equation for *Nematostella* and the exact analytical solution for *Exaiptasia*. We will then analyze these solutions and demonstrate that predatory stinging increases with starvation, whereas defensive stinging stays constant or decreases. These results do not require specific definitions of the parameters but hold under the broad conditions discussed below.

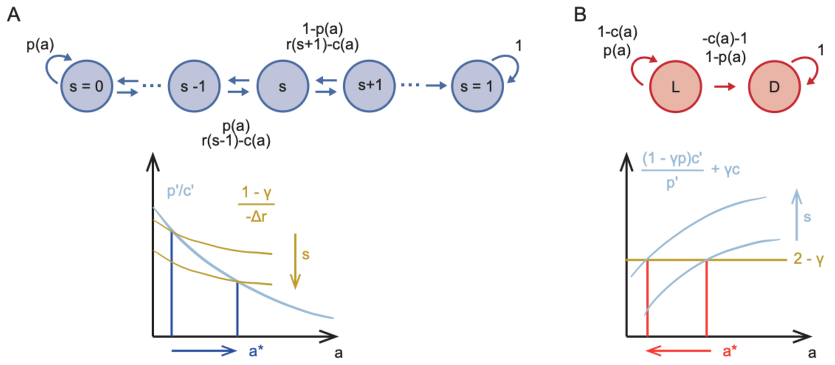


Supplementary Figure 1: Sketch and results of Markov Decision Process for predatory stinging modeling *Nematostella* behavior (A) and for defensive stinging modeling *Exaiptasia* behavior (B).

#### Asymptotic solution for optimal predatory stinging

In the interest of clarity, we repeat the setup of the model and some basic notions in control theory. We consider a Markov decision process, where an agent is in a state of starvation $s$ that goes from $0$ (well fed) to $1$ (maximally starved, as sketched in Supplementary Figure 1A, top). Transitions occur at a sequence of time points where the agent is presented with a prey and can either catch it and transition to better satiation $s \to s' = s - 1$ (with probability $p(a)$) or miss it and transition to $s \to s' = s+1$ (with probability $1-p(a)$). The action or control $a$ represents the intensity of the sting, and goes from $0$ (no stinging) to $1$ (all nematocytes discharged at once); $p(a)$ thus grows with $a$ as discharging more nematocysts increases chances to successfully envenomate the prey. The agent acts according to a policy $a \sim \pi(s)$, which prescribes the action that is played at each state $s$ (in general $\pi(s)$ is a probability distribution that maps states to actions). We introduce the immediate reward function (which we call "desirability" in the main text) and assume that satiation entails higher rewards, hence $r(s')$ is a decreasing function of $s'$. Additionally, because nematocytes are single-use cells, we assume their discharge has a cost $c(a)$ increasing with $a$, hence the net reward is $r(s') - c(a)$.

The goal of control theory is to find the optimal policy of actions that allows maximizing the net cumulative reward in the future:

$$\pi^*(s) = \arg\max_{\pi} \left[ E \left( \sum_{t=0}^{\infty} (\gamma^t r_{t+1}(s_t, a_t, s_{t+1}) | s_0 = s \right) \right]$$

where $s_t$ is the state at time $t$, $a_t \sim \pi(s_t)$ is the action chosen in the current state, $s_{t+1}$ is the state the agent transitions to as prescribed by the transition rates and $r_{t+1}$ is the immediate reward obtained upon transitioning. As usual, because we have an infinite horizon, we consider discounted rewards where $\gamma < 1$; for smaller and smaller values of $\gamma$ the long-term reward is less and less important in making current decisions. To derive the optimal policy we define the quality factor matrix

$Q^{\pi}(a,s)$ which measures the expected cumulative reward given that the agent starts in state $s$ and plays action $a$ and then continues with a policy $a \sim \pi(s)$

$$Q^{\pi}(a,s) = E\left(\sum_{t=0}^{\infty}(\gamma^k r_{t+1}(s_k, a_k, s_{k+1})|s_0 = s, a_0 = a\right)$$

At optimality, i.e. for the optimal policy $\pi^*(s)$, the quality factor obeys a fixed point equation known as Bellman's optimality equation:

$$Q(a,s) = \sum_{s'} p(s'|a,s)[r(s,a,s') + \gamma \max_{a'} Q(a', s')]$$

where $p(s'|a,s)$ are the transition probabilities and $Q(a,s)$ is the maximum expected sum of all future net rewards starting from the starvation state $s$ and taking action $a$. The optimal policy is deterministic and satisfies:

$$a^*(s) = \arg\max_a Q(a,s)$$

It is often useful to introduce the value function $V^*(s)$, corresponding to the maximum expected cumulative reward starting from state $s$:

$$V^*(s) = \max_a Q(a,s) \tag{1}$$

so that

$$Q(a,s) = \sum_{s'} p(s'|a,s)[r(s,a,s') + \gamma V^*(s)]$$

In our problem, only transitions to the two adjacent states are allowed, hence the sums in Equation (1) simplify in:

$$\begin{aligned} V^*(s) &= \max_a\{p(a)[r(s-1) - c(a) + \gamma V^*(s-1)] + (1-p(a))[r(s+1) - c(a) + \gamma V^*(s+1)]\} \\ &= \max_a\{-c(a) + p(a)[r(s-1) + \gamma V^*(s-1)] + (1-(p(a))[r(s+1) + \gamma V^*(s+1)]\} \quad (2) \end{aligned}$$

*Short hand notation*: $a^* \equiv a^*(s)$.
By zeroing the derivative we obtain the maximizer $a^*$ (when $a^* \in (0,1)$):

$$-c'(a^*) + p'(a^*)[r(s-1) + \gamma V^*(s-1)] - p'(a^*)[r(s+1) + \gamma V^*(s+1)] = 0$$

$$[r(s-1) + \gamma V^*(s-1)] = r(s+1) + \gamma V^*(s+1) + \frac{c'(a^*)}{p'(a^*)} \tag{3}$$

Plugging Equation (3) into the Bellman Equation (2) we obtain:

$$V^*(s) = -c(a^*) + r(s+1) + \gamma V^*(s+1) + p(a^*)\frac{c'(a^*)}{p'(a^*)} \tag{4}$$

From Equation (3) there is also

$$V^*(s-1) = V^*(s+1) + \frac{1}{\gamma}[r(s+1) - r(s-1)] + \frac{c'(a^*)}{p'(a^*)} \tag{5}$$

Equations (4), (5) are two equations in the four unknowns $V^*(s)$, $V^*(s+1)$, $V^*(s-1)$ and $a^*(s)$. These equations can be solved iteratively by coupling all states and using the boundary conditions on the absorbing state. However, the exact iterative solution is not particularly illuminating.

Instead, we will make a simplifying assumption that leads to a good approximation, which is much simpler to handle.
Assume that $a^*(s)$ varies slowly with $s$ so that $a^*(s) \approx a^*(s+1) \approx a^*(s-1)$ (better approximations may be achieved by assuming a first order expansion). Then we obtain a third equation by writing Equation (4) for the state $\bar{s} = s - 1$

$$V^*(s-1) = -c(a^*) + r(s) + \gamma V^*(s) + p(a^*)\frac{c'(a^*)}{p'(a^*)} \tag{6}$$

We can then repeat the trick to obtain a fourth equation. To this end, we first eliminate $V^*(s-1)$ by combining Equations (6) and using also (5) we obtain:

$$V^*(s+1) = K(a^*)\frac{\gamma}{1-\gamma^2} - \frac{r(s+1)}{\gamma} + \frac{r(s-1)}{\gamma(1-\gamma^2)} + \frac{r(s)}{1-\gamma^2} \tag{7}$$

$$K(a^*) = -c(a^*)\frac{1+\gamma}{\gamma} + \frac{c'(a^*)}{p'(a^*)}\left(p(a^*)\frac{1+\gamma}{\gamma} - \frac{1}{\gamma^2}\right) \tag{8}$$

Repeating the trick, we can write Equation (7) for $\bar{s} = s - 1$, and using that $a^*(s-1) = a^*(s)$ and obtain a fourth equation to close the system:

$$V^*(s) = K(a^*)\frac{\gamma}{1-\gamma^2} - \frac{r(s)}{\gamma} + \frac{r(s-2)}{\gamma(1-\gamma^2)} + \frac{r(s-1)}{1-\gamma^2} \tag{9}$$

The system is now closed with 4 equations in the 4 unknowns. We solve for $a^*$ by eliminating $V^*(s)$ from Equations (4) and (9) and plugging Equation (7) for $V^*(s+1)$ as a function of $a^*$. After some (tedious) algebra, we obtain that $a^*$ satisfies the simple relation:

$$\frac{p'(a^*)}{c'(a^*)} = \frac{1-\gamma}{-\Delta r(s)} \tag{10}$$

Both the asymptotic solution from Equation (10) and the numerical solution from value iteration are used in the main text (main text Figure 2B right, symbols and lines respectively). Match of asymptotics and numerics occurs over a wide variety of functional forms of the reward function and parameters (see Figure 4). The asymptotics break down if abrupt changes in the rewards and transition rates are assumed, which leads to exceeding slopes in the optimal policy (data not shown). Equation (10) has a non-trivial solution $0 < a^* < 1$ when $c' > 0$, $r(s)$ is a decreasing function, and $p(a)$ is an increasing function. If we additionally assume that $r$ is concave, and $p'/c'$ is a decreasing function of $a$ (for example, $p$ is strictly concave and $c$ is convex), then Equation (10) prescribes that $a^*$ increases with $s$, as seen graphically in Figure 1A, bottom. Hence independently of the specific functional forms of $c$, $p$, and $r$, as long as these broad assumptions are valid, optimal stinging for predation entails more intense attacks as starvation increases. For different assumptions of reward function $r(s)$, cost function $c(a)$, and probability $p(a)$, we can easily substitute the specific expressions into Equation (10) and solve for $a^*$ for every $s$.

When the cost function depends on both state and action $c = c_0(s)a$, and specifically if $c_0(s)$ increases with starvation, comparisons between numerics and asymptotics suggest that the asymptotic solution in equation (10) still holds (data not shown). From the asymptotic solution, it is easy to visualize why predatory stinging grows with starvation even if the cost increases (moderately) with starvation. Indeed, if $c$ increases slightly with $s$, the light blue curve in Supplementary Figure 2 slightly shifts downward with $s$. If the shift is sufficiently small, its intersection with the green curves still occurs for increasing values of $a$ (dashed line in Supplementary Figure 2). However, a dramatic increase of $c$ with $s$ will shift the blue curve downward considerably, and the intersection will eventually move backward (dotted line in Supplementary Figure 2). In other words, when the cost of nematocyst discharge for starved animals is dramatically larger than for well-fed animals, the benefits of predation are outweighed by its cost and the optimal predatory stinging decreases with starvation (exemplified in Supplementary Figure 3B, green and yellow curves). Note that such a dramatic increase in the cost with starvation may be unrealistic, as it entails that the cost of starvation for well-fed animals is comparatively negligible (see green and yellow cost functions in Supplementary Figure 3A). A formal proof of the asymptotic solution for this case and further consequences for Markov Decision Processes on a continuous state space are beyond the scope of the current paper and are subject of ongoing work.

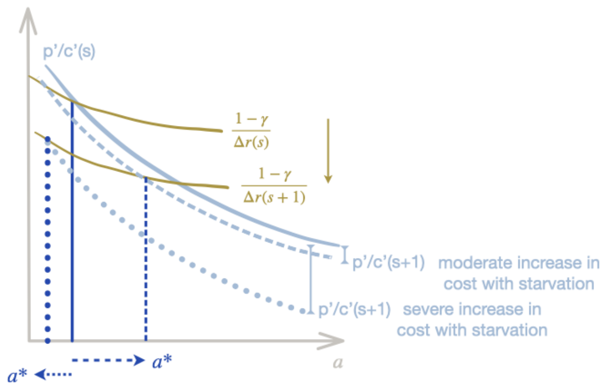


Supplementary Figure 2: Same as Supplementary Figure 1(A), but for a cost per nematocyte that varies with starvation $c = c_0(s)a$. Moderate increases in the cost per starvation do not affect the qualitative results. More dramatic increases do lead to a decrease in predatory stinging with a starvation state.

#### Analytical solution for optimal defensive stinging

To model defensive stinging, we assume the agent is in a safe state $L$ and attacks for defense against external threats (see the sketch in Supplementary Figure 1B, top). An attack with intensity $a$ leads to successful defense hence remaining in the same state $L$ with probability $p(a)$, it entails a cost $c(a)$, and unit reward. The same attack can lead to failed defense and a transition to the state of danger $D$ with probability $1 - p(a)$. $D$ is absorbing and it entails a penalty of $-1$. Hence Bellman optimality equation for the sole non-trivial state $L$ is

$$V^*(L) = \max_a\{(1-p(a))(-1-c(a)) + p(a)(-c(a) + \gamma V^*(L) + 1)\} = \tag{11}$$

$$= \max_a\{-c(a) - (1-p(a)) + p(a)(\gamma V^*(L) + 1)\} \tag{12}$$

Let $\arg\max_a V^*(L) = a^*$ ; if $a^* \in (0, 1)$, then

$$-c'(a^*) + p'(a^*)[1 + \gamma V^*(L) + 1] = 0 \tag{13}$$

Combining Equation 13 with Equation 12, we can cancel off $V^*$ and obtain an equation for $a^*$ only:

$$(\gamma p(a^*) - 1)\left(\frac{c'(a^*)}{p'(a^*)} - 1\right) = \gamma c(a^*) + \gamma(1 - p(a^*)) - 1 \tag{14}$$

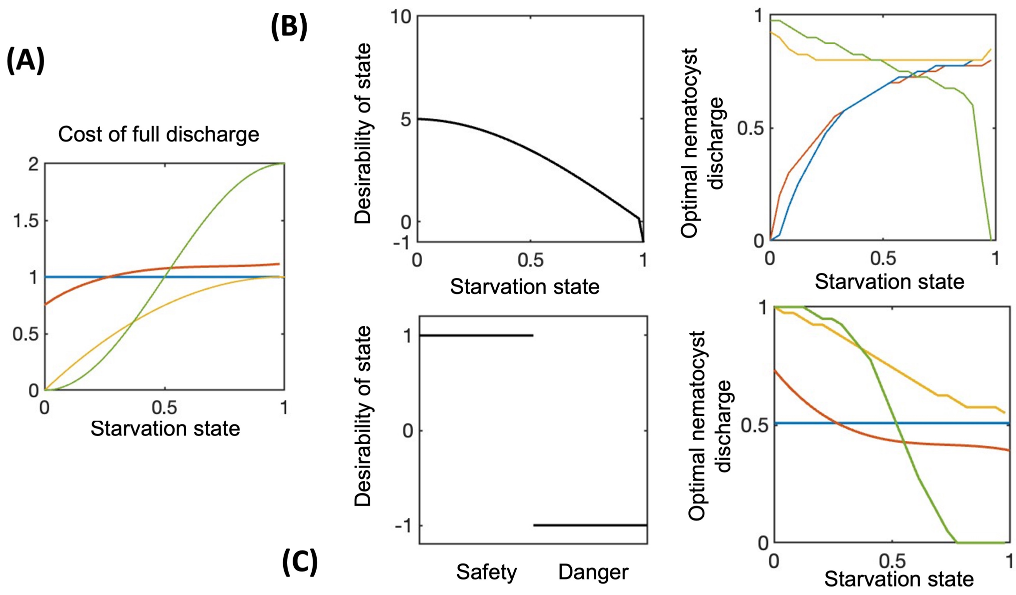


Supplementary Figure 3: Predictions for predatory and defensive stinging with different cost functions; compare with Figure 2B in the main text. For a cost of full discharge that stays constant or increases moderately with starvation (panel A, blue and red line respectively), the predicted optimal stinging is discussed in the main text. Here we consider two additional cost functions that increase more dramatically with starvation (panel A, yellow and green lines). For *Nematostella* (panel B), we define reward as in the main text desirability 2 Figure 2B; the new cost functions cause a decreasing optimal nematocyst discharge (corresponding yellow and green lines in panel B, right, numerical solutions). For *Exaptasia* (panel C), the more dramatically decreasing cost functions still result in decreasing optimal discharge (C, right, numerical solutions).

Equation (14) prescribes a constant optimal strategy $a^*$, which value depends on the details of the assumptions for $p$ and $c$.
Thus for the same assumptions used for the previous case, optimal defensive stinging does not depend on starvation. To further elaborate on this point, we note that although the state of satiation of the agent is not modeled, we may consider that whatever state of starvation the agent happens to be in will modulate the cost of stinging. For example, shooting at maximum discharge may be more metabolically costly if the animal is starving. In this case, Equation (14) will depend parametrically on $s$ through the cost $c = c_s(a)$:

$$(\gamma p(a^*) - 1)\left(\frac{c_s'(a^*)}{p'(a^*)} - 1\right) = \gamma c_s(a^*) + \gamma(1 - p(a^*)) - 1 \quad (15)$$

where $c'$ and $c''$ are derivatives with respect to $a$. For specific assumptions of the cost function $c_s(a)$, and probability $p(a)$, we can easily substitute the expressions into Equation (15) and solve for $a^*$ for every $s$. For example, if $c = c_0(s)a$ and $p = p_M a(2-a)$, the solution is $a^* = K - \sqrt{K^2 - C}$ where $K = \epsilon/c_0(s)$; $\epsilon = (2-\gamma)/\gamma$, $C = -1/p_M\gamma + 2\epsilon/c_0(s)$ and it exists for $c_0(s) < 2p_M(2-\gamma)$. Outside these boundaries, the solution is either $a^* = 0$ or $a^* = 1$, and it is obtained by comparing $V(L)$ for these two choices of action and choosing the one that maximizes $V$. This solution is decreasing as can be easily demonstrated by deriving with respect to $s$. We used this solution to fit the experimental data and obtained the cost function shown in the main text (main text Figure2

A bottom, empty circles).

The optimal stinging strategy $a^*$ decreases with $s$ in much more general conditions e.g. using the same broad assumptions discussed in the previous section for predatory stinging: $p(a)$ is concave and $c$ is convex in $a$ (either $p$ or $c$ can be linear in $a$, but not both). Additionally, we assume that $c$ does not decrease with starvation, e.g. $c = c_0(s)a$ where $c_0' \geq 0$. Under these conditions, the solution $a^*$ decreases or remains constant with $s$. To prove this, we simply note that $a^*$ is defined by the point where:

$$\gamma \frac{1-\gamma p}{p'} c_s' + \gamma c_s = 2 - \gamma \tag{16}$$

If the l.h.s. of Equation (16) is an increasing function of $a$, and if $c_s(a)$ increases with $s$, the intersection of the l.h.s. with the constant value $2-\gamma$ is found graphically as in Figure 1B, bottom and occurs at lower and lower values of $a$ as $s$ increases. The l.h.s. of Equation (16) is indeed an increasing function of $a$ as seen by deriving with respect to $a$ and using the assumptions: $p < 1$; $p'' \leq 0$; $c' \geq 0$ and $c'' \geq 0$.

Thus under the same broad assumptions for the functional forms of $c$ and $p$, stinging for predation increases with starvation, whereas stinging for defense remains constant or decreases with starvation.

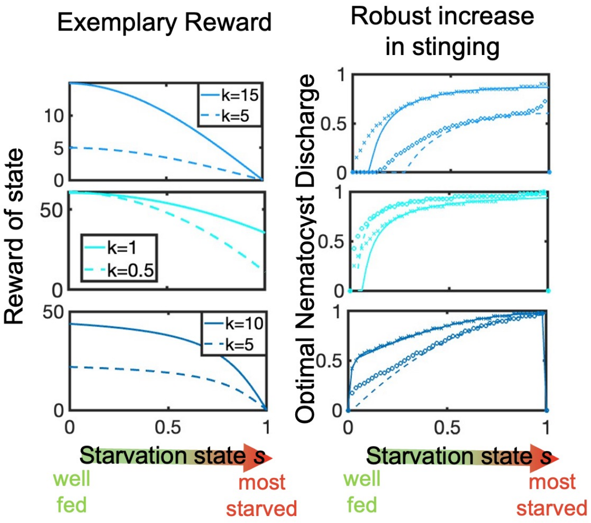


Supplementary Figure 4: Optimal policy predicted by Bellman's equation for the MDP sketched in Supplementary Figure 1A. Left: three choices of concave reward functions $r(s')$ ($r(s) = k\cos(s\pi/2)$, upper left; $r = k(1-50s^2)+60$, middle left; $r = k\tan^{-1}(5(1-s)/(\pi/10))$, lower left); solid and dashed lines correspond to two choices of the parameter $k$ for each reward as in the legend. The cost of full dischare is $c_0 = 1.5$ and the likelihood of successful discharge is as in the main text with $p_M = 0.6$. Right: the asymptotic solution for the optimal policy $a^*(s)$ (solid and dashed lines matching the corresponding reward on the left) reproduces well the numerical solution obtain from numerically solving Bellman's Equation (1) (crosses and circles correspond to the solid and dashed rewards on the left). Optimal nematocyst discharge increases with the starvation state independently on the shape of the reward function.
